# DeORFanizing *Candida albicans* Genes using Co-Expression

**DOI:** 10.1101/2020.12.04.412718

**Authors:** Teresa R. O’Meara, Matthew J. O’Meara

**Author notes:** Corresponding Authors: Teresa R. O’Meara, Department of Microbiology and Immunology University of Michigan Medical School, 1150 W. Medical Center Dr, Medical Sciences Building II, Room 6751 Ann Arbor, MI, 48109, USA, Matthew J. O’Meara, Department of Computational Medicine and Bioinformatics University of Michigan Medical School, 1150 W. Medical Center Dr, Palmer Commons, 100 Washtenaw Ave #2017, Ann Arbor, MI 48109, USA.

## Abstract

Functional characterization of open reading frames in non-model organisms, such as the common opportunistic fungal pathogen *Candida albicans*, can be labor intensive. To meet this challenge, we built a comprehensive and unbiased co-expression network for *C. albicans*, which we call CalCEN, from data collected from 853 RNA sequencing runs from 18 large scale studies deposited in the NCBI Sequence Read Archive. Retrospectively, CalCEN is highly predictive of known gene function annotations and can be synergistically combined with sequence similarity and interaction networks in *Saccharomyces cerevisiae* through orthology for additional accuracy in gene function prediction. To prospectively demonstrate the utility of the co-expression network in *C. albicans*, we predicted the function of under-annotated open reading frames (ORF)s and identified *CCJ1* as a novel cell cycle regulator in *C. albicans*. This study provides a tool for future systems biology analyses of gene function in *C. albicans.* We provide a computational pipeline for building and analyzing the co-expression network and CalCEN itself at (http://github.com/momeara/CalCEN).

**Importance:** *Candida albicans* is a common and deadly fungal pathogen of humans, yet the genome of this organism contains many genes of unknown function. By determining gene function, we can help identify essential genes, new virulence factors, or new regulators of drug resistance, and thereby give new targets for antifungal development. Here, we use information from large scale RNAseq studies and generate a *C. albicans* co-expression network (CalCEN) that is robust and able to predict gene function. We demonstrate the utility of this network in both retrospective and prospective testing, and use CalCEN to predict a role for C4_06590W/*CCJ1* in cell cycle. This tool will allow for a better characterization of under-annotated genes in pathogenic yeasts.

## Introduction

Co-expression analysis is based on the hypothesis that genes that are coordinately expressed under multiple diverse conditions and perturbations are likely to function in the same biological process (1). Co-expression networks are built from transcriptomic studies across a range of conditions, incorporating broad and unbiased analyses of gene expression at a global scale. The gene-by-condition matrix found in most transcriptomic studies can be used for differential gene expression analysis, where the effects of a perturbation or a mutation can be compared to a background or wildtype. In contrast, a co-expression network transforms the gene-by-condition matrix into a gene-by-gene matrix, or equivalently, a gene network, where the edge weight defines the degree of co-expression. To estimate this co-expression, several methods have been proposed, including the Pearson correlation coefficient, the Spearman correlation coefficient, or a partial correlation coefficient (2). Although co-expression approaches to identify gene function have been used extensively in the model yeast *S. cerevisiae* and humans (1, 3, 4), it has only recently been applied to full effect in other fungi, as in the recent work from Meyer and colleagues (5, 6). Not only does co-expression have high predictive accuracy for gene function annotations, it also captures evolutionary-scale changes in cell identity (7). For non-model organisms such as *Candida albicans*, two important questions are 1) is there utility in building a species-specific co-expression network, or are all the relevant co-expression signatures available through orthology with a model organism (e.g., *S. cerevisiae*); and 2) how extensive does the transcriptional profiling need to be to generate useful co-expression networks? The second question is particularly relevant for emerging infectious diseases, as broad transcriptional profiling takes significant research investment to generate.

*Candida albicans* is a common opportunistic pathogen of humans; it is both an asymptomatic colonizer of the mucosal surface of healthy individuals and a deadly invasive pathogen in immunocompromised patients. There is significant effort in determining gene function for *C. albicans,* with the goal of identifying essential genes, virulence factors, or regulators of drug resistance and thus expanding the target space for future antifungal development. Gene Ontology (GO) term annotation is one framework for describing gene function. For a gene to be fully annotated, it requires an understanding of the molecular function (MF), biological process (BP), and the cellular component (CC). The *C. albicans* genome consists of 6,468 ORFs, and approximately 70% of these remain under-characterized (8). As an extreme set, 1,801 ORFs have no Biological Process GO term annotation in the Candida Genome Database (9). In many cases, gene annotation information comes from inferred orthology with the model yeast *S. cerevisiae,* but these organisms diverged approximately 150 MYA (10), and even gene essentiality is not necessarily conserved across these species (8, 11). Current high-throughput approaches for gene function analysis include transposon insertion screens (11, 12), functional genomic screens of mutant libraries (8, 13–16), genetic interaction screens (17), yeast-two-hybrid approaches (18), and other protein-protein interaction screens (19). However, there are also many transcriptomic analyses of *C. albicans* available through NCBI SRA, suggesting that it is now feasible to create a co-expression network for *C. albicans* as a complementary approach for predicting gene function.

Here, we generated a robust co-expression network from 853 sequencing runs from 18 available transcriptomic datasets for *C. albicans,* using rank-correlation through the EGAD R package to build the network. We then added information from other modalities, including sequence similarity and *S. cerevisiae* protein-protein interactions from BIOGRID, as incorporation of orthogonal information allows for better gene function prediction (20). Retrospective analysis of the clustering identified high network connectivity of histone proteins and ribosomal proteins, validating the efficacy of this approach. We also demonstrate that there are distinct sub-networks for different GPI-anchored cell wall proteins. We then applied this co-expression network to examining genes of unknown function in *C. albicans* and identified Ccj1 as a DnaJ-containing protein that acts as a regulator of cell cycle. This co-expression resource can be used by the research community for examination of gene function and network connectivity in *C. albicans.*

## Results

### Co-expression networks in *Candida albicans*

A co-expression network is built by through three stages: 1) collecting transcriptome data over multiple environmental conditions to generate a gene-by-condition expression matrix, 2) measuring the correlation between all pairs of gene expression profiles to generate a gene-by-gene correlation matrix, and 3) interpreting the correlation matrix as a network where the nodes represent genes and the edge weights represent the degree of co-expression. To build the *<u>C</u>andida <u>a</u>lbicans* <u>c</u>o-<u>e</u>xpression <u>n</u>etwork (CalCEN), we identified RNA sequencing (RNAseq) studies from the NCBI Sequence Read Archives (SRA), which we then filtered for studies with at least 20 *C. albicans* samples based on the guidelines from Ballouz and colleagues (20–22), yielding 12 unpaired and 6 paired end studies, listed in Supplemental Table 1. The conditions for these experiments included differences in carbon source, co-culture with host cells or bacteria, treatment with chemical perturbations, or differences in mutations, highlighting the diversity in conditions covered by these studies. To ensure that all studies were processed consistently, we collected the raw reads from SRA and re-aligned all of the data to the *C. albicans* SC5314 genome Assembly 22 coding transcripts using RSEM with bowtie2 (21–23) and generated a heatmap combining of all of the RNAseq reads (Figure 1A). Since RNAseq of co-cultured samples can lead to low coverage depth, we removed runs where greater than 50% percent of the genes had zero expression (Supplemental Figure 1), yielding 853 runs in total. To control for the bias that more reads will map to longer genes, we used the fragments per kilobase of transcript per million mapped reads (FPKM) as the estimated expression for each gene under each condition. Because the primary aim of our study is to evaluate the utility of co-expression as a data source, we used the simple yet robust Spearman-rank correlation to measure the correlation between gene expression profiles, as implemented in the EGAD R package (24) (Figure 1B). Thus, for each pair of genes, we have a value between 0 and 1 representing the rank of co-expression among all pairs of genes.

**Figure 1:**
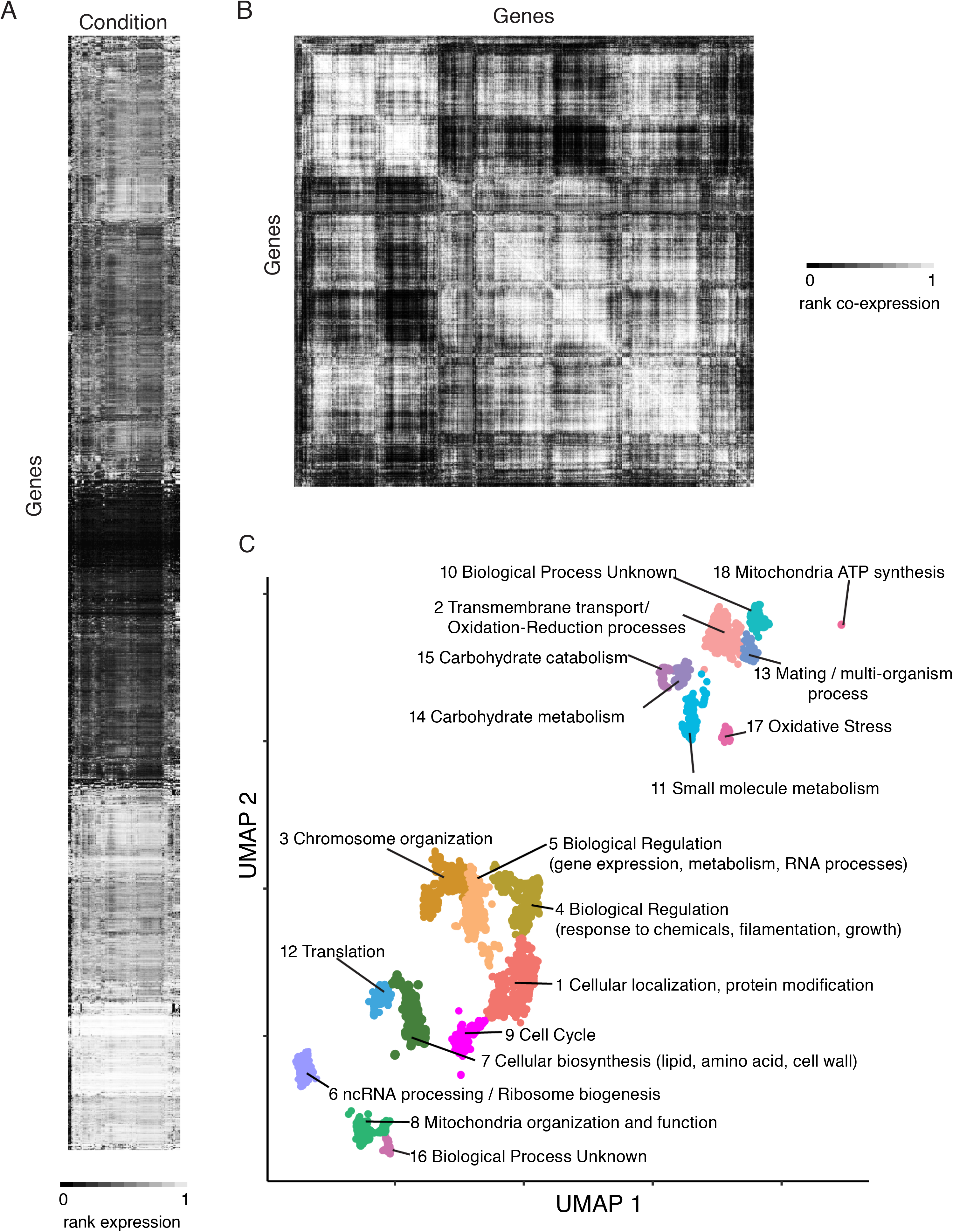
Generating a co-expression network for *C. albicans*. A) A gene-by-environment heatmap generated from collected *C. albicans* RNAseq experiments from the SRA. The *C. albicans* genes are on the Y-axis and conditions are on the X-axis. B) A gene-by-gene heatmap generated from Spearman rank correlation. C) UMAP embedding reveals functional clusters. Annotations were determined by GO term enrichment of genes in each cluster.

To visualize the co-expression network, we projected the network to two dimensions using UMAP (25), which aims to keep co-expressed genes closer together than non-co-expressed genes. Interestingly, we identify 18 clusters distinct clusters (Supplemental Table 2). Using Gene Set Enrichment Analysis (26) and GO term enrichment of the genes in each cluster, we can observe clear functional signatures (Figure 1C), including cluster 9 which is enriched for cell cycle proteins, or cluster 18, which is enriched for proteins encoded on the mitochondrial genome.

### Predicting gene function using multiple modalities

There are many networks that can be built for predicting gene function, and each can provide information about currently under-annotated genes, and the different modalities can be combined for greater coverage and accuracy (20). To measure the information in CalCEN that is not already captured by other modalities, we compared the co-expression network to the BlastP, SacGene, SacPhys, and YeastNet genome-scale networks. Filtering the co-expression network for the top 1% of co-expressed genes shows substantial coverage of the information captured in other networks, while also having information on over 550 genes that are not included in any other network (Figure 2A).

**Figure 2:**
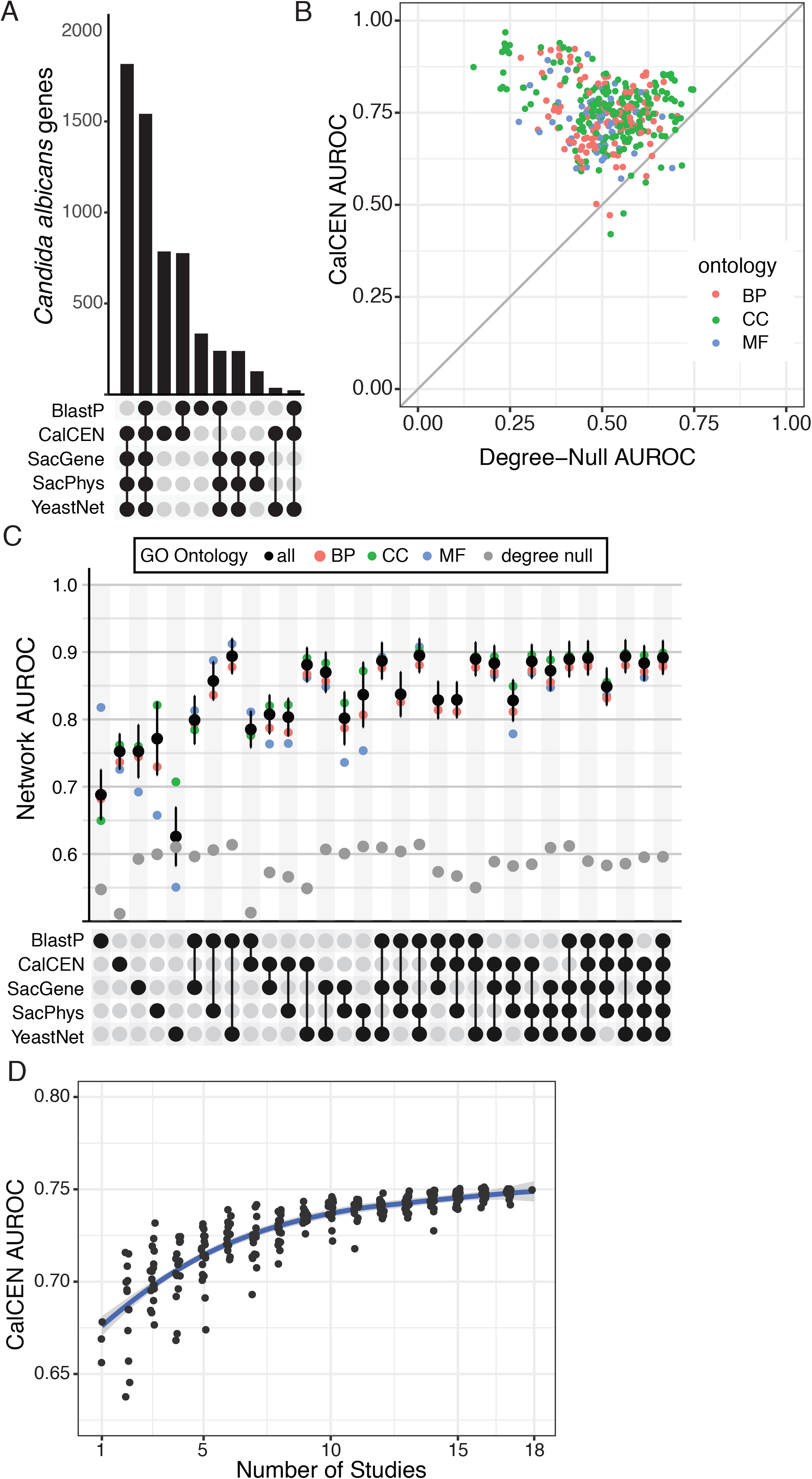
CalCEN provides a robust orthogonal approach to identifying gene function. **A)** UpSet plot of network overlap. BlastP network is thresholded at Evalue < 1e-5, CalCEN thresholded at top 1%. Each bar in the upper region shows the number of gene-nodes in the intersection of the set of networks indicated by the rows with filled circles in the lower region. **B)** For each annotated GO term colored by ontology biological process (BP), cellular component (CC) or molecular function (MF), the CalCEN neighbor-voting guilt-by-association (GBA) area under the ROC curve (AUROC) is plotted as a function of the degree-null (genes predicted based on their network degree) AUROC. **C)** Mean neighbor-voting GBA performance for individual and combined networks, indicted by the rows with filled circles in the lower region, for sub-ontology terms (colored dots), all terms (black) with error bars representing the standard error in the mean over 10-fold cross validation replicas. Degree-null predictive accuracy for each network is shown in grey. **D)** Mean neighbor-voting GBA performance for the CalCEN built over random subsets of RNAseq studies. The blue curve represents a mean of a non-parametric LOESS fit with standard deviation in dark grey. As the number of studies increases, the performance increases.

To benchmark the utility of our co-expression network to predict gene function, we first performed a retrospective prediction of GO term annotations collected from the Candida Genome Database and FungiDB (9, 27). Guilt by Association (GBA) is a method to predict gene function by propagating annotation labels through a given network. We implement GBA prediction through neighbor voting, where the strength of a term predicted for a gene is determined by the fraction of neighbors in the network having the term. For each term, we constructed a ranked list of genes by the strength of the prediction and compared it to the known set of annotations using the area under the receiver operating characteristic curve (AUROC) score and averaging over a 10-fold cross validation.

We began by measuring the predictive accuracy of the network relative to uninformative baselines. The null baseline predictor that randomly predicts genes for each term yields an AUROC of on average 0.5. However, a stronger but also non-informative baseline for a network sorts the genes by their number of neighbors and predicts this single sorted list for every term. This degree null predictor is often a surprisingly strong predictor for retrospective gene function prediction because multifunctional genes, such as well studied signaling hubs, tend to be both well connected and annotated for many functions. However, this predictor has low utility for prospective gene function prediction as it cannot find new genes for a function of interest, and it cannot find a new function for a gene of interest. If a network has a high degree null predictor relative to its neighbor voting predictor, it suggests that the neighbor voting predictor may be biased towards multifunctional genes (28, 29). The BlastP network, derived from whole gene sequence similarity (BlastP) has an average degree null AUROC of 0.55 ± 0.045 and network AUROC of 0.69 ± .037. Orthology networks from *S. cerevisiae*, derived from physical protein-protein interactions (SacPhys), genetic interactions (SacGene) curated from BioGRID have average degree null AUROCs of 0.60 ± 0.042 and 0.59 ± 0.042 and average network AUROCs of 0.78 ± 0.054 and 0.75 ± 0.039. The integrated *S. cerevisiae* network (YeastNet) that includes which incorporates literature co-annotations has an average degree null AUROC of 0.61 ± 0.046 and an average network AUROC of 0.63 ± 0.044. We find that the CalCEN network has an average degree null AUROC of 0.51 ± 0.034 and a neighbor voting AUROC of 0.75 ± 0.026, indicating that the network has low multifunctionality bias relative to other networks (Figure 2B, C and Supplemental Figure 2).

When comparing the predictive accuracy of co-expression (Co-Exp) to the other networks, we see that the Co-Exp has network has predictivity comparable to SacGene and SacPhys, and significantly better than BlastP and YeastNet. When the networks are combined additively, we see that adding Co-Exp improves the predictive accuracy of each other network, with Co-Exp and YeastNet being particularly predictive (Figure 2C). Together, this demonstrates that we are capturing information with the CalCEN that is not included in previous networks, and that addition of the CalCEN can improve gene function predictions.

A challenge in building the CalCEN was determining whether the number of RNAseq studies was sufficient to generate a robust network. To assess this, we asked how the predictive accuracy of the CalCEN changes based on the number of RNA-seq studies used for generating the network. Embedding each RNAseq run based on the expression pattern for each gene (Supplemental Figure 3), we see that the runs cluster by study, suggesting the between-study condition differences are greater than the within-study condition differences, and that additional studies would increase coverage of the expression topology. To assess the impact of study diversity on retrospective gene function prediction, we sampled subsets of studies and re-computed the network and GBA performance. We found that the mean performance increased from ~.675 to .75 from 1 to 18 studies, suggesting that a minimum of 10 RNAseq studies is needed for competitive performance compared with previous predictive methods, but that even at 18 studies, the performance of the co-expression network for GBA predictions has not yet saturated and may be improved when additional studies are added.

### Retrospective Identification of Conserved Gene Clusters

We then examined our co-expression network for its ability to identify specific gene clusters that have been previously identified in other organisms. Previous work in *S. cerevisiae* described a co-expression cluster of ribosomal proteins and other proteins involved in translation, as these proteins are coordinately regulated in response to nutrient conditions (1, 4). We used a set of known ribosomal proteins to seed the network, identified all of the first neighbors in the network, and then identified the co-expression edges for all of the genes in this set. This resulted in a densely connected cluster for many known ribosomal and proteasomal proteins in *C. albicans* (Figure 3A). However, we also identified some known ribosomal proteins that were not contained within the main cluster, suggesting potentially differential regulation patterns.

**Figure 3:**
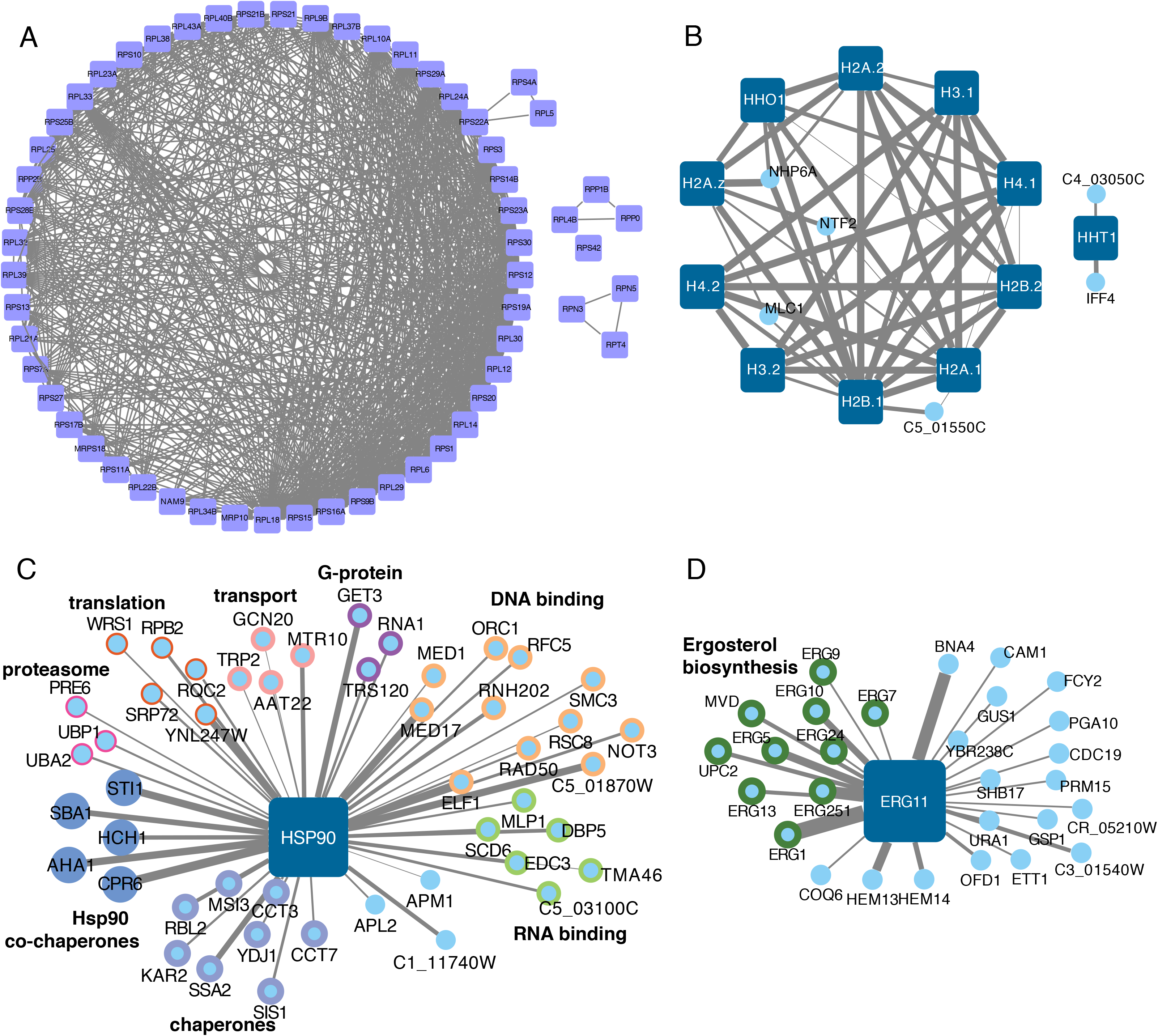
Retrospective analysis identifies functional clusters of genes in *C. albicans.* A) Ribosomal proteins form a densely connected co-expression cluster. B) Histone proteins are highly connected, except for the HHT1 variant histone protein. C) Hsp90 is co-expressed with Hsp90 co-chaperones, as well as multiple other functional classes of proteins. D) Erg11 is co-expressed with other components of the ergosterol biosynthetic cascade. Nodes represent the genes, and the edge width corresponds to the degree of co-expression.

Another well-known cluster of proteins are the histone proteins, which are transcriptionally regulated with the cell cycle. In *S. cerevisiae,* this cluster is composed of 8 histone proteins (1), but in *C. albicans,* we observed that 10 histone proteins cluster with each other (Figure 3B). Both H2A variant proteins (H2A.1 and H2A.2) are present in the *C. albicans* histone cluster cluster (30); however, the H3 variant gene, *H3.A / HHT1*, is not connected with this cluster. This is consistent with the recent reports of a decreased abundance of *HHT1* compared with the canonical H3 proteins *HHT2* and *HHT21,* and the co-expression of *HHT1* with *IFF4* and *CPS1* is consistent with a potential transcriptional connection between *HHT1* and the biofilm circuit described by Rai et al. (31). In the main histone cluster, we also observed *NTF2,* a Nuclear envelope protein, and *NHP6A,* a High-mobility group (HMG) protein that binds to and remodels nucleosomes, as connected with multiple histone proteins. Intriguingly, C5_01550C, a protein of unknown function, was also connected with both H2B.1 and H2A.1; however, this protein is also co-expressed with ribosomal proteins.

Hsp90 is a conserved and essential molecular chaperone that physically interacts with many *C. albicans* proteins to regulate their folding and function (19). The co-expression network for Hsp90 was able to identify 5 Hsp90 co-chaperones (Figure 3C) and 7 additional chaperone proteins. However, it also identified clusters of genes involved in many core aspects of cell biology, including protein translation and degradation, consistent with the pleiotropic role of Hsp90 in the cell. In addition to proteins that act in a complex, we hypothesized that our network would identify genes that act in a single biosynthetic cascade. To test this, we examined our network for genes that co-express with Erg11, the major target of the azole antifungals and part of the ergosterol biosynthetic cascade (32). By using just Erg11 as the seed, we identified 8 additional genes in the ergosterol biosynthetic cascade that were co-expressed with Erg11 (Figure 3D). Moreover, we identified Upc2, the transcription factor that regulates ergosterol biosynthesis, as part of this cluster. Notably, this approach also identified multiple genes involved in heme uptake, as well as others that are involved in general metabolism.

### GPI-anchored proteins form multiple, distinct co-expression clusters

In *C. albicans,* the fungal cell wall plays an essential role in regulating interactions with host cells and tissues (33– 36). The outer layer of the cell wall has many glycosylated proteins that are anchored into the cell wall via GPI motifs (37), and many of these proteins are not fully characterized. We selected a set of 27 GPI-anchored proteins as seeds and examined their co-expression networks (Figure 4A). This revealed that while there was a subset of GPI-anchored proteins that cluster with known cell wall biogenesis proteins, such as Pga38 and Pga54, there were also distinct and non-overlapping co-expression networks that did not include cell wall related processes. For example, Pga10 was most associated with proteins involved in metabolism, such as the Eno1 enolase, the Cdc19 pyruvate kinase, and the Pfk1 and Pfk2 phosphofructokinases, consistent with the role of Pga10 in using heme and hemoglobin as iron sources (38). Pga63, which is proposed to be a component of the COPII vesicle coat, is co-expressed with other secretion proteins (Sec61, Sec23, Sec24) and proteins involved in N-glycosylation. In contrast, Pga27, which is currently un-annotated, is co-expressed with multiple transcription factors and the Rim21 and Sln1 signal transduction proteins that regulate cell wall remodeling in response to stress, which suggests that Pga27 may play a role in sensing or responding to stress. Similarly, the unannotated protein Pga59 is co-expressed with multiple proteins involved in RNA metabolism, including tRNA synthetases, while Pga57 is co-expressed with mitochondrial proteins. This highlights the potentially diverse biological functions of GPI-anchored proteins in *C. albicans* biology.

**Figure 4:**
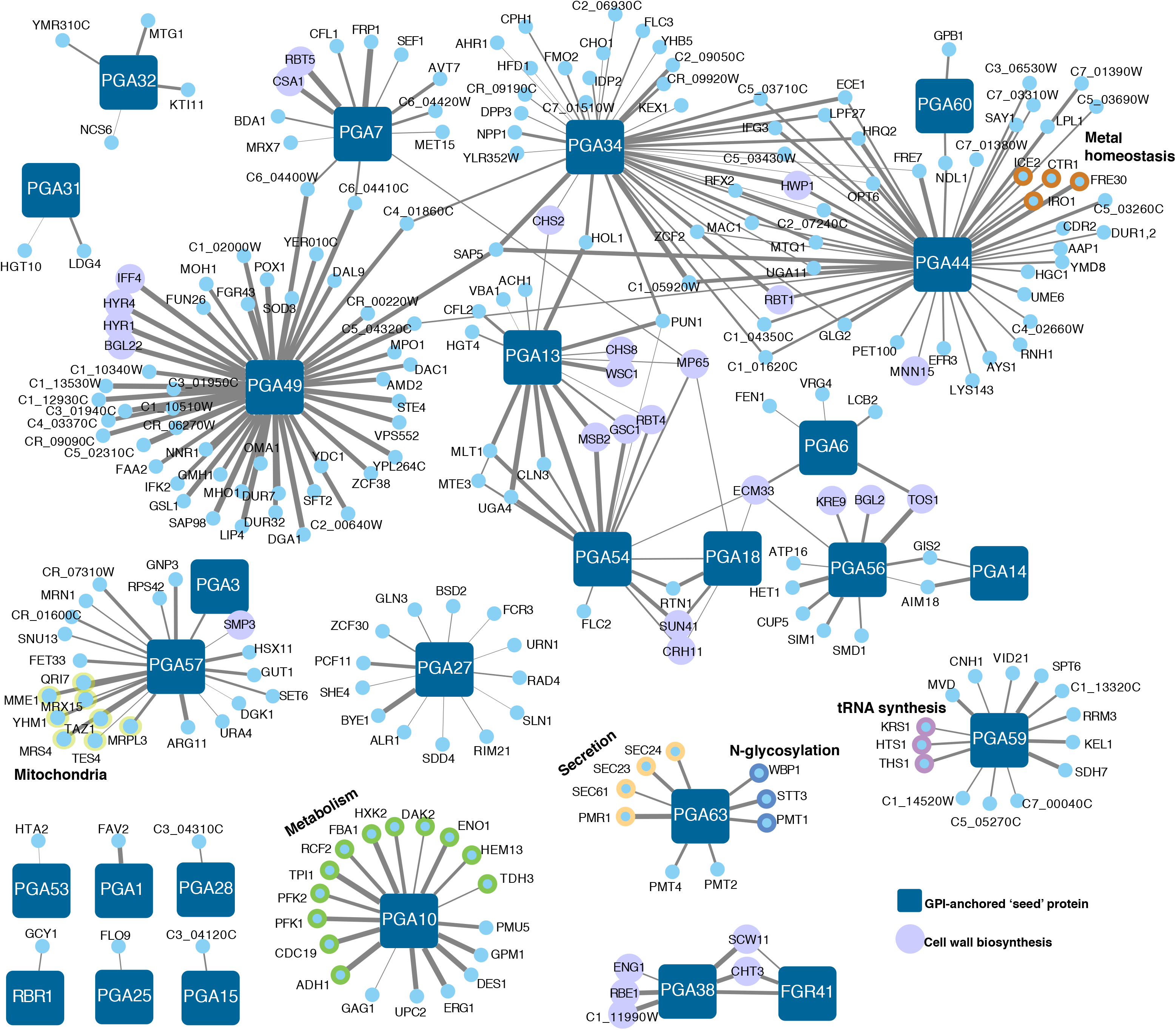
GPI-anchored proteins in *C. albicans* show distinct co-expression clusters. 27 GPI-anchored proteins were used as seeds to generate co-expression clusters. Genes were included as co-expressed if the passed the top 2% cutoff.

### Prospective testing of the co-expression network identifies a new role in cell cycle regulation for uncharacterized gene C4_06590W/*CCJ1*

Many genes in the *C. albicans* genome do not have an assigned GO term annotation. The co-expression network can provide insight into genes without current annotation, suggesting functions that can then be tested experimentally. The gene C4_06590W is an uncharacterized protein that is present in the CTG clade but not in the model yeast *S. cerevisiae.* We generated a model for the N-termini of the C4_06590W structure using TrRosetta (39) and compared it with the crystal structure of the Sis1 DnaJ-containing protein from *S. cerevisiae* (40), which highlighted the conservation of the DnaJ domain (Figure 5A). To investigate the function of C4_06590W, we examined the co-expression network using C4_06590W as the seed protein. This identified multiple genes involved in *C. albicans* cell cycle as clustering with C4_06590W (Figure 5B), which led us to hypothesize that C4_06590W may be involved in cell cycle. In *C. albicans,* mutants in cell cycle often result in aberrant filamentation in the absence of an inducing cue (8, 41–44). Therefore, we examined the phenotype of the *tetO-C4_06590W/C4_06590W* mutant strain in the presence or absence of doxycycline to repress target gene transcription (Figure 5B). When C4_06590W was repressed, the majority of the cells showed aberrant germ tube formation (Figure 5C). However, nuclear division appeared normal, with each cell containing just one nuclei (Figure 5C). We then tested growth in the presence of hydroxyurea, a DNA synthesis inhibitor that induces S-phase arrest (44). Depletion of C4_06590W resulted in hypersensitivity to hydroxyurea, as observed by a decrease in growth (Figure 5D) and a striking increase in the number of filaments at subinhibitory concentrations of the drug (Figure 5E). Together, these data suggest that C4_06590W is involved in cell cycle control in *C. albicans,* and we have therefore proposed *CCJ1*, for Cell Cycle DnaJ, as the new gene name for C4_06590W.

**Figure 5:**
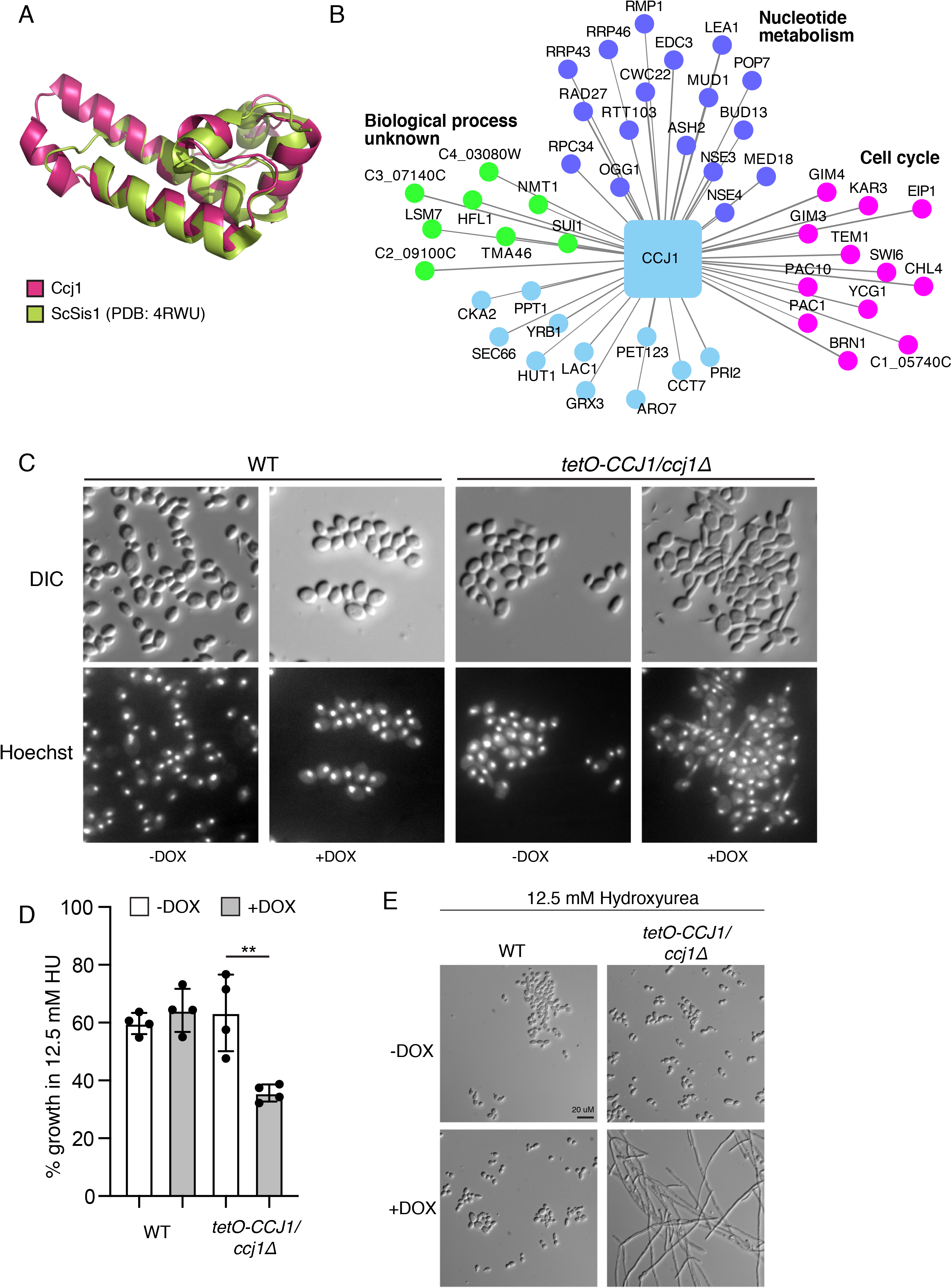
Identification of C4_06590W, or *CCJ1,* as a novel cell cycle regulator. A) Ccj1 contains a J domain. TrRosetta was used to generate a de novo fold for the Ccj1 protein, and the structure was overlaid on the *S. cerevisiae* Sis1 DnaJ protein crystal structure. B) The co-expression network for Ccj1 identifies multiple cell cycle proteins. C) Repression of *CCJ1* with 5 μg/mL DOX results in aberrant filamentation in the absence of an inducing cue. Nuclei were stained using Hoechst. D) Repression of *CCJ1* with 5 μg/mL DOX results in hypersensitivity to 12.5 mM hydroxyurea. Data from two biological replicates, with two technical replicates each. E) Repression of Ccj1 results in hyperfilamentation in response to 12.5 mM hydroxyurea. Cells were incubated in YPD with 12.5 mM hydroxyurea and the presence or absence of 5 μg/mL of doxycycline overnight before imaging.

## Discussion

Co-expression has been shown to be a reliable means of identifying genes that share function in a variety of contexts. Genome scale co-expression networks built from bulk RNAseq studies have been used extensively for gene function prediction using guilt by association (1, 2, 45). Recently, co-expression networks have been built for non-model organisms, including *Aspergillus* (5, 6). Here, we develop the CalCEN, a co-expression network for *C. albicans* from available large-scale, publicly available RNAseq studies. We show that this the network is built from sufficient data to meaningfully organize genes into functional modules and retrospectively predict gene annotations. We demonstrate the utility of this network to predict functional networks around several classes of genes, including ribosomal proteins and the ergosterol biosynthetic cascade. Moreover, the CalCEN analysis of histone proteins accurately identified a separation of the non-canonical Hht1 histone H3 protein (31) from the canonical histone protein complex, whereas both H2A variants (30) are highly co-expressed. Co-expression analysis of GPI-anchored proteins in *C. albicans* showed a variety of clusters with different enrichments for biological functions, highlighting that not all cell wall associated proteins are involved in cell wall biosynthesis. Additionally, we used the CalCEN to identify a new regulator of cell cycle, *CCJ1*, which we verify contains a DnaJ domain using *de novo* structure prediction. We propose that the CalCEN will provide a method for predicting gene function for the *C. albicans* research community.

A limitation of this study is that we use simplistic methods for generating and analyzing the network. For example, we used Spearman rank correlation for building the network, we defined arbitrary thresholds when needed (such as the top 1% to compare network overlap), we combined networks by adding network weights, performed guilt by association using neighbor voting, and evaluated retrospective enrichment by using the area under the ROC curve. While more sophisticated methods may give higher predictive accuracy, an advantage of these simplistic methods is that they transparently show how the data lead to the predictions, without complex global normalization. Further, the predictive accuracy of the simple methods gives a lower bound on the utility of the data. Thus, in demonstrating the utility of the CalCEN, and making the methods and networks available, we hope to enable their wider use in the fungal pathogenesis community, where the methods can be refined and integrated with additional data sources for gene function prediction.

## Methods

### Network Construction

**<u>CalCEN</u>**: RNA expression was estimated by aligning reads to *Candida albicans* SC5314, Assembly 22 (46) coding transcripts by converting from SRA to FASTQ format using fastq-dump from the NCBI SRA tools package and then aligned using the RSEM package v1.2.31(47) with bowtie2 (47) using the default settings. Reads that did not map or mapped multiple times were discarded, and the percent that mapped exactly once are shown in (Figure S1), plotted using the ggplot2 package (48) for R. The CalCEN network over genes was estimated by the Spearman-Rank correlation coefficients of the FPKM values across all runs using the EGAD R package (45). **<u>BlastP</u>**: A sequence similarity network was created comparing all pairs of protein transcripts from *Candida albicans* SC5314, Assembly 22 using Protein-Protein BLAST 2.2.30+ (51), yielding 103,400 associations. Protein-protein interaction networks were created from data collected from BioGRID build (3.4.161) (48) for, yielding 110,991 orthologous physical interactions between 3,953 *Candida albicans* genes (**<u>SacPhys</u>**) and 395,437 orthologous genetic interactions between 3942 *C. albicans* genes (**<u>SacGene</u>**). These protein protein networks were then extended to include indirect associations with weights inverse of the shortest path (52). **<u>YeastNet</u>** v3, which is an integrative network for *S. cerevisiae* built from co-citation, co-expression, co-occurrence of protein domains, genomic context, genetic interactions, high and low throughput protein-protein interactions, phylogenetic profiles between yeast genes and 3D structure of interacting orthologues (49). The overlap of the genes and interactions of these networks is shown in (Figure 2) using the UpSetR R package (50).

To assess the biological relevance of the co-expression network, functional annotations for curated by the Candida Genome Database (http://www.candidagenome.org/download/go/gene_association.cgd.gz) were collected on 12-Jun-2018, filtering for terms with qualifier != “NOT” and propagating up “is_a” and “part_of” term relationships using the GO.db R package (51). Terms were then filtered for those having at least 20 and at most 1000 annotations, yielding 9,144 annotations for 169 biological process (BP) terms, 19,672 annotations for 215 cellular component (CC) terms, and 4,741 annotations for 85 molecular function (MF) terms. By evidence ~73% annotations were inferred from electronic annotation (IEA), ~11% were inferred from a mutant phenotype (IMP), ~8% were inferred from direct assay (IDA) and less than one percent annotations for other evidence codes.

### Embedding of the Co-Exp network

To embed the CalCEN genes, we used Monocle 3 (52) to preprocess the 6226 x 853 expression matrix using principal component analysis from 853 dimensions to 100 dimensions and then applied the UMAP algorithm to reduce the dimensions from 100 to 2. UMAP was used with parameters a = 50, b = 0.5, n_neighbors = 30, n_epochs = 2000, negative_sample_rate = 50, and repulsion_strength = 3, which builds on the R implementation of UMAP, uwot, (54) https://github.com/jlmelville/uwot), and the Approximate Nearest Neighbors Oh Yeah library (https://github.com/spotify/annoy). Clusters were identified using Leiden community detection algorithm (53) with parameters k = 30, num_iter = 10, resolution = 0.1 and plotted with ggplot2 (54).

### Gene set construction

6468 *Candida albicans* open reading frames were identified in the SC5314 Assembly 22 (21) database and downloaded from FungiDB (54) (Supplemental Table 3). 1801 ORFs in this set did not contain a computed or curated GO term annotation for Biological Process. 27 GPI-anchored proteins were identified in the Candida Genome Database. Ribosomal proteins were identified by curated GO term annotation, and the first neighbors of the seed network were identified using the CalCEN.

### Protein structure prediction for CCJ1

To predict the structure of the CCJ1 DnaJ domain, the sequence for residues 9-96 were submitted to the TrRosetta de novo structure prediction server (38), which built a deep multiple sequence alignment of 19,612 sequences and used a machine learning model to estimate the distances and relative angles of each pair of residues. These contact maps were then used to sample coarse-grain and full-atom protein folding conformation spaces to optimize a molecular mechanics forcefield to find low-energy conformations. The top resulting conformations were highly consistent with pairwise full atom RMSD 0.15 Å, and when structurally aligned with 4RWU, a 1.25 Å X-ray crystal structure of the DnaJ containing protein Sis1 from in *Saccharomyces cerevisiae*, it has a full-atom RMSD of 1.6 Å (Figure 5A), confirming the Interpro and Pfam sequence-based DnaJ domain annotation.

### CalCEN computational workflow

Computational methods implemented here are available as an R package (github.com/momeara/CalCEN).

### Network Visualization

All network visualizations were generated in Cytoscape (55).

### Strains, reagents, and culture conditions

All *C. albicans* strains were archived in 25% glycerol and stored at −80°C. Overnight cultures were grown in YPD (1% yeast extract, 2% bacto peptone, 2% dextrose) at 30°C with rotation. The *tetO-C4_06590W/C4_06590W* strain and the associated parental wild type strain used in this study were created as part of the GRACE tetracycline-repressible mutant collection (8, 56). Doxycycline (Fisher MP219504410) was dissolved in water and used at the indicated concentrations. To repress gene expression, overnight cultures of the relevant strains and controls were subcultured in the presence or absence of 5 μg/mL DOX before use. Hydroxyurea (ThermoFisher A10831-06) was dissolved in water and used at the indicated concentrations. Hoechst (Cayman Chemical 15547) was dissolved in water and used at 1 μg/mL.

### MIC assays

Drug tolerance assays were performed in flat bottom 96-well plates (Alkali Scientific) using a modified broth microdilution protocol, as previously described (19). The assays were performed in a total volume of 0.2 mL/well with 2-fold dilutions of the drug in YPD. Plates were incubated in the dark at 30C for 24 hours before reading OD600 values on a spectrophotometer (Synergy H1). Each strain was tested in technical and biological duplicate. Data at a single concentration of drug (the highest concentration where growth of the wild type strain was not inhibited) is displayed.

### Microscopy

To monitor *C. albicans* morphology, we used differential interference contrast (DIC) microscopy on an Olympus iX70 inverted microscope and a Hamamatsu FLASH4 CMOS camera at 40X magnification. For fluorescence microscopy, we used an X-cite series 120 light source with a DAPI filter set. To monitor nuclei, cells were fixed and permeabilized with methanol before addition of 1 ug/mL Hoechst and imaging. Representative images from two biological replicates are shown.

## Supplemental Information

Table S1: RNA-Seq studies for *Candida albicans* co-expression analysis

Table S2: Co-expression clusters from UMAP

**Figure S1:**
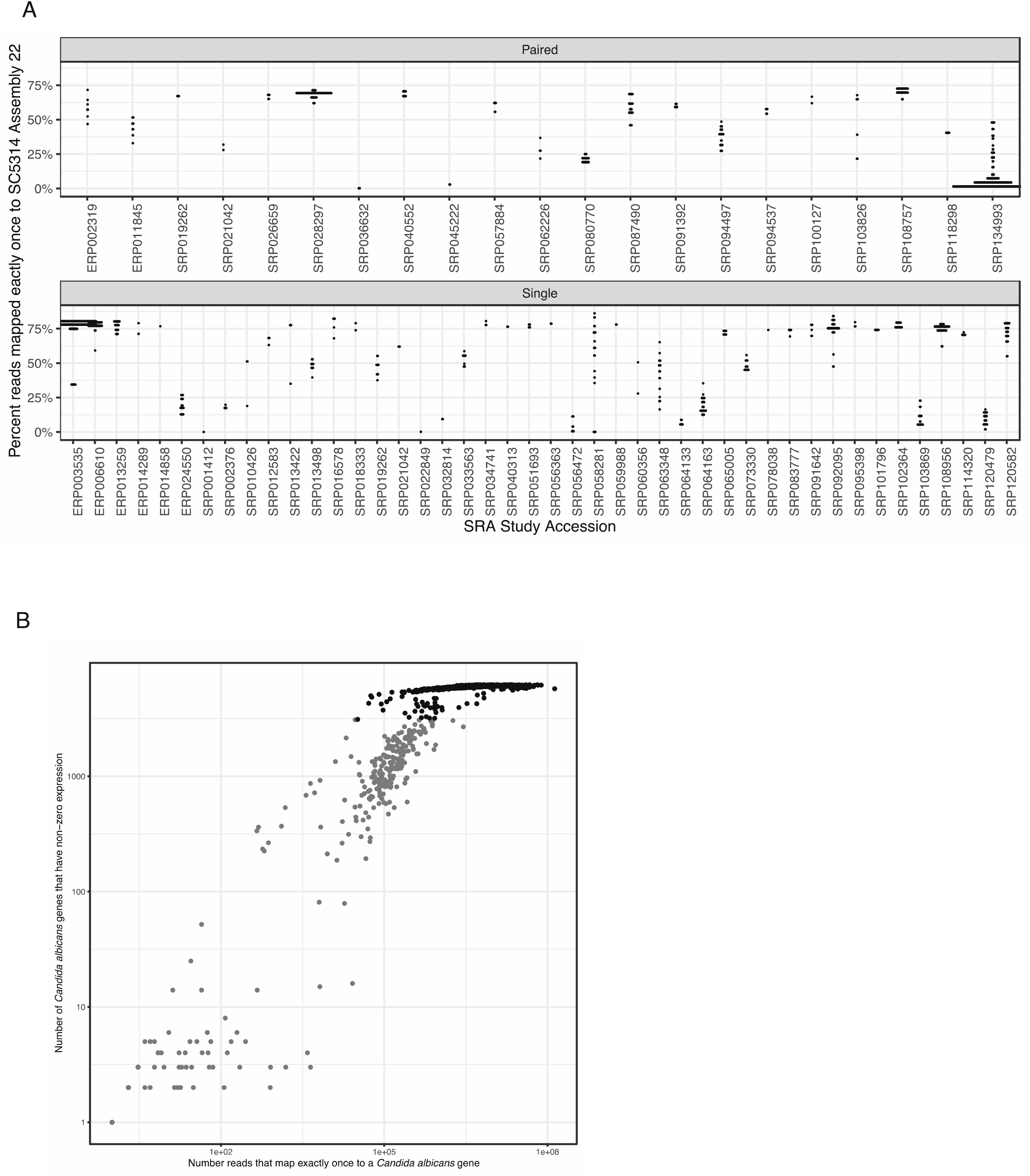
**A)** 1399 RNA-seq runs from 18 identified studies are scatter-plotted as the number of genes with non-zero expression vs. the fraction transcripts that map exactly once. **B)** The 853 runs that have non-zero expression for at least half of the genes (3113), which are used to construct the CalCEN network, are shown in black.

**Figure S2:**
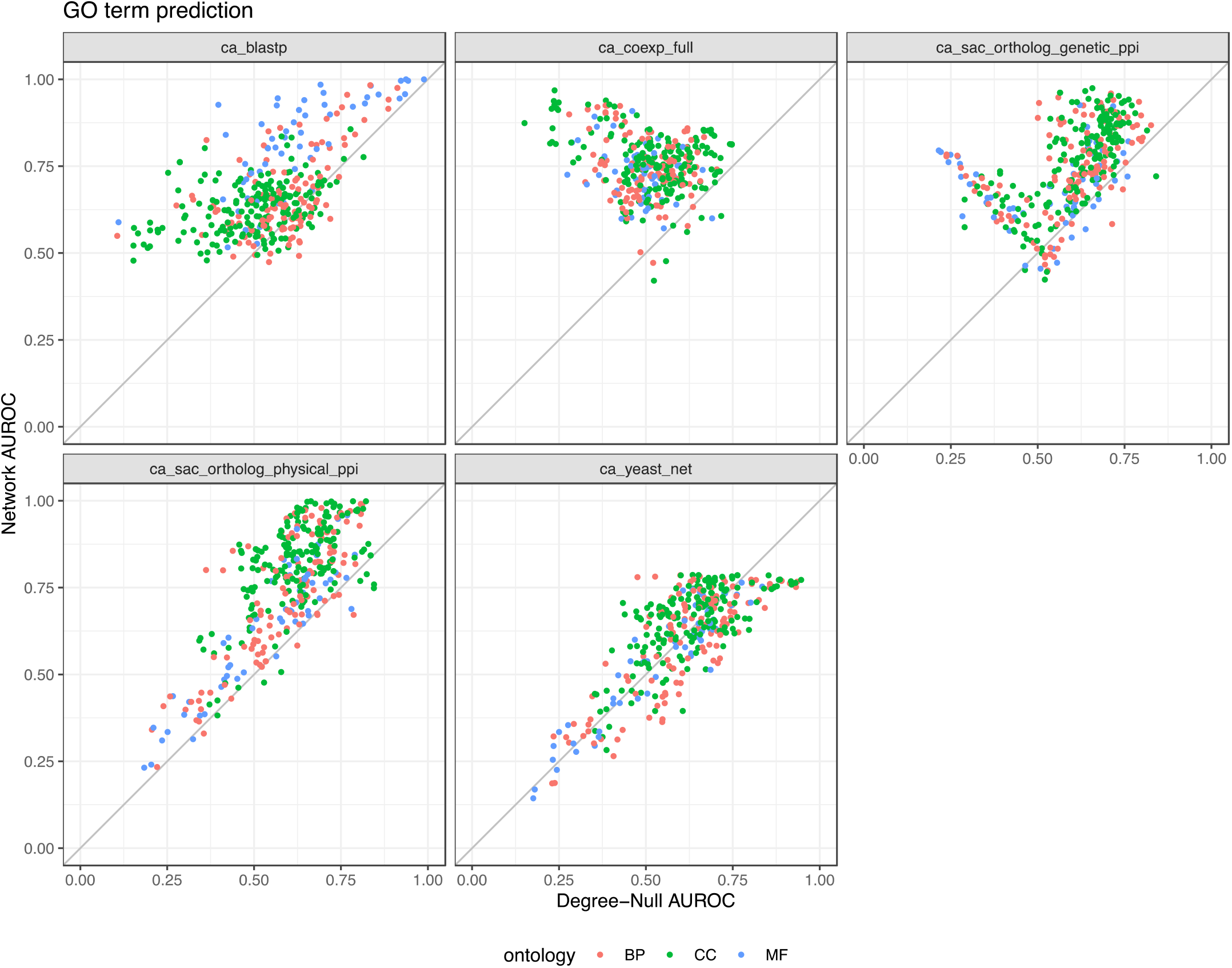
For each annotated GO term colored by ontology biological process (BP), cellular component (CC) or molecular function (MF), the indicated network neighbor-voting guilt-by-association (GBA) area under the ROC curve (AUROC) is plotted as a function of the degree-null (genes predicted based on their network degree) AUROC.

**Figure S3:**
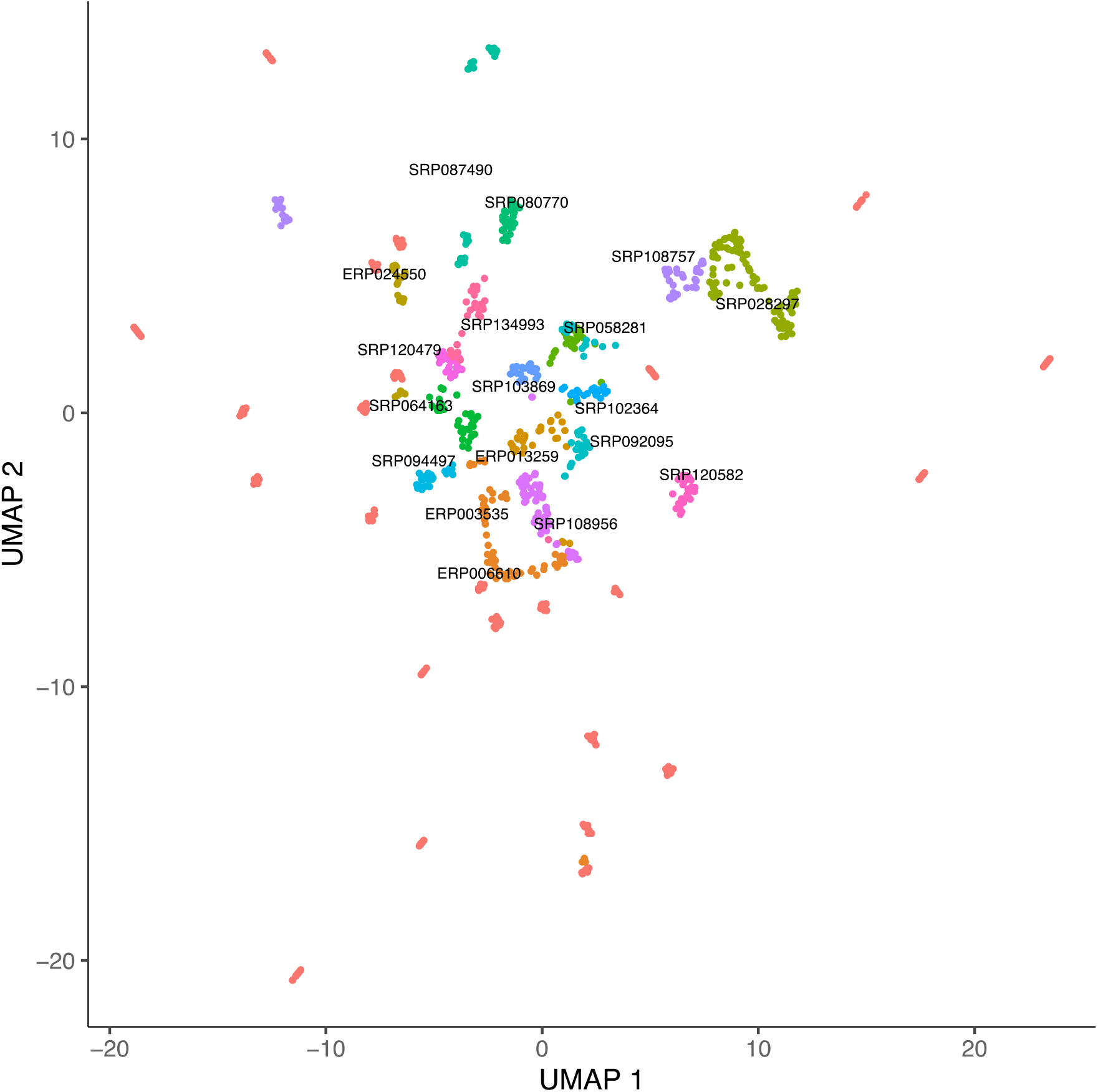
The 853 RNA-seq runs are embedded by reducing the 6226 dimension gene-expression profiles to 100 dimensions using principal component analysis and then to 2 dimensions using UMAP using min_dist=0.5 and n_neighbors=30. Runs were then labeled by their study accession and plotted using ggplot2.

Networks: CalCEN, BlastP, SacGene, SacPhys, YeastNet

## Acknowledgements

This research was supported in part through computational resources and services provided by Advanced Research Computing at the University of Michigan, Ann Arbor. TRO was supported by the University of Michigan Biological Sciences Scholars Program (BSSP). We thank Merck and the University of Toronto for making the GRACE *C. albicans* mutant collection available.

